# The psychotomimetic ketamine disrupts the transfer of late sensory information in the corticothalamic network

**DOI:** 10.1101/2022.02.21.476564

**Authors:** Yi Qin, Ali Mahdavi, Marine Bertschy, Paul M Anderson, Sofya Kulikova, Didier Pinault

## Abstract

In prodromal and early schizophrenia, disorders of attention and perception are associated with structural and chemical brain abnormalities, and with dysfunctional corticothalamic networks exhibiting disturbed brain rhythms. The underlying mechanisms are elusive. The non-competitive NMDA receptor antagonist ketamine simulates the symptoms of prodromal and early schizophrenia, including disturbances in ongoing and task & sensory-related broadband beta-/gamma-frequency (17-29 Hz/30-80 Hz) oscillations in corticothalamic networks. In normal healthy subjects and rodents, complex integration processes, like sensory perception, induce transient, large-scale synchronized beta/gamma oscillations in a time window of a few hundreds of ms (200-700 ms) after the presentation of the object of attention (e.g., sensory stimulation). Our goal was to use an electrophysiological multisite network approach to investigate, in lightly anesthetized rats, the effects of a single psychotomimetic dose (2.5 mg/kg, subcutaneous) of ketamine on sensory stimulus-induced oscillations. Ketamine transiently increased the power of baseline beta/gamma oscillations and decreased sensory-induced beta/gamma oscillations. In addition, it disrupted information transferability in both the somatosensory thalamus and the related cortex and decreased the sensory-induced thalamocortical connectivity in the broadband gamma range. In conclusion, the present findings support the hypothesis that NMDA receptor antagonism disrupts the transfer of perceptual information in the somatosensory cortico-thalamo-cortical system.

**LAY ABSTRACT:** Cognitive deficit is usual in schizophrenia. Perception- or task-related beta/gamma-frequency oscillations are decreased. In healthy humans and rodents, ketamine-induced NMDA receptor antagonism simulates the symptoms of early schizophrenia and excessively amplifies baseline beta/gamma oscillations. In the present study, using an electrophysiological multisite network approach in a rodent model, it is demonstrated that ketamine, systemically administered at a single psychotomimetic dose, increases baseline beta/gamma oscillations, decreases beta/gamma responses induced by sensory stimulation in a short time window (200-700 ms), and disrupts information transfer in the cortico-thalamo-cortical network. The present findings have mechanistic relevance for cognitive deficits in schizophrenia.

## INTRODUCTION

In many neuropsychiatric illnesses, including schizophrenia, sleep disorders and deficits in attention-related sensorimotor and cognitive integration processes are common. These disorders insidiously start to occur during the prodromal phase (McGhie & Chapman, 1961; Lunsford-Avery *et al*., 2013; Manoach *et al*., 2014; Zanini *et al*., 2015; Mayeli *et al*., 2021). Administration of the non-competitive NMDA receptor antagonist ketamine at a subanesthetic dose can, after the administration of a subanesthetic dose, induce a psychosis-relevant mental state in healthy humans (Krystal *et al*., 1994; Hetem *et al*., 2000; Anticevic *et al*., 2015; Hoflich *et al*., 2015; Rivolta *et al*., 2015; Grent-’t-Jong *et al*., 2018) and other species, including rodents (Chrobak *et al*., 2008; Pinault, 2008; Pitsikas *et al*., 2008; Ehrlichman *et al*., 2009; Hakami *et al*., 2009; Kocsis, 2012). The ketamine-induced psychosis-relevant mental state is reminiscent of both the prodromal phase of schizophrenia and the psychotic transition.

Sensory-related perception is a very complex and relatively long-lasting (∼ 2 s) process, which involves early (<200 ms) and late (>200 ms) stages. These two-time stages represent a continuum through highly distributed systems involving diverse cortical areas during the perceptual process (Portella *et al*., 2012; Portella *et al*., 2014; Saradjian *et al*., 2019). The dynamics of the cortico-thalamo-cortical (CTC) network in the late stage of perception remains little known in psychotic disorders. Literature suggests the existence of a link between late sensory-related activities and perception. In a visual perception task, participants show sensory perception-related gamma activity increases in two separate components, early and late (Rodriguez *et al*., 1999). Likewise, in a study conducted in humans and mice two response activities (early:< 300 ms, and late: > 300 ms) are recorded after visual stimulation, the latter response believed to be involved in visual perception (Funayama *et al*., 2015).

In schizophrenia patients, deficits in perception are associated with a reduction of phase synchrony in beta/gamma-frequency (20-60 Hz) oscillations at the late period (Uhlhaas *et al*., 2013). These results suggest a decrease in induced sensory-related gamma oscillations during the late period of perception. There are lines of evidence showing a decrease in induced-gamma oscillations in individuals with a clinically at-risk mental state for psychotic transition (Haenschel *et al*., 2009; Reilly *et al*., 2018). The decrease in power and synchrony of task & sensory-induced gamma oscillations may be due to the abnormal amplification of basal gamma oscillations recorded in such patients (Ramyead *et al*., 2015). Indeed, in the acute rodent ketamine model, early sensory-evoked gamma oscillations decrease whereas ongoing gamma oscillations increase, supporting the hypothesis of a reduction in the signal-to-noise ratio (Hakami *et al*., 2009; Kulikova *et al*., 2012; Anderson *et al*., 2017).

We wanted to study whether and how late sensory-induced beta/gamma oscillations (17-29 Hz/30-80 Hz) are disturbed by NMDA receptor antagonism. To do so, we investigated the effects of low-dose ketamine in the somatosensory thalamocortical (TC) system during the late sensory stimulus-related period (200-700 ms post-sensory stimulation), involving highly distributed CTC systems (Alitto & Usrey, 2003; Briggs & Usrey, 2008; Urbain *et al*., 2015; Homma *et al*., 2017). Since sensory-induced gamma oscillations can be recorded in anesthetized rats (Neville & Haberly, 2003), the experiments were conducted in the pentobarbital-sedated rat. Spectral analysis and coherence connectivity were used in an attempt to estimate, respectively, the level of synchronization and the functional connectivity between the recording sites (Kam *et al*., 2013). Unlike amplitude measures, coherence measurement shows the synchronization level between two signals based on the phase consistency (Srinivasan *et al*., 2007). EEG and extracellular signals are relatively complex as they are generated by multiple interacting cortical and subcortical oscillators. The complexity of such signals, related to functional aspects of the corresponding neural networks, can be assessed with non-linear analyses such as the multiscale entropy analysis (MSE) (Costa *et al*., 2005; Miskovic *et al*., 2019). The MSE has been applied to EEG from psychiatric patients (Fernandez *et al*., 2013). Higher MSE can indicate increases in the complexity of time-varying signals and may represent disruptions in long-range temporal connectivity or temporal integration (Breakspear & Stam, 2005). So, in an attempt to measure the dynamical complexity in the TC system at multiple timescales, MSE was applied to the extracellular local field potential (LFP) recordings. The present findings show that, in the somatosensory CTC system, ketamine disrupts the information transfer of sensory-induced gamma oscillations.

## MATERIAL AND METHODS

### Animals and drugs

Seven adult (3-6-month-old, 285-370 g), Wistar male rats were used. All animal care procedures were performed under the approval of the Ministère de l’Education Nationale, de l’Enseignement Supérieur et de la Recherche. Animals were housed and kept under controlled environmental conditions (temperature: 22 ± 1°C; humidity: 55 ± 10%; 12 h/12 h light/dark cycle; lights on at 7:00 am) with food and water available *ad libitum*. Every precaution was taken to minimize stress and the number of animals used for each series of experiments. Ketamine (Imalgene 1000, MERIAL), pentobarbital sodique (CEVA santé animale) and fentanyl (Fentadon@ DECHRA) were from CENTRAVET.

### Surgery under deep narco-analgesia

Narcosis was initiated with an intraperitoneal injection of pentobarbital (60 mg/kg). An additional dose (10–15 mg/kg) was administered as soon as there was a nociceptive reflex. Analgesia was achieved with a subcutaneous injection of fentanyl (7.5 μg/kg) every 30 min. The depth of the surgical narco-analgesia was continuously monitored using an electrocardiogram, watching the rhythm and breathing, and assessing the nociceptive withdrawal reflex. The rectal temperature was maintained at 36.5°C (peroperative and protective hypothermia) using a thermoregulated pad. The trachea was cannulated and connected to a ventilator (50% air–50% O^2^, 60 breaths/min). Under local anesthesia (lidocaine), an incision of the skin on the skull was done, and the periosteum was removed to set the skullcap bared and to perform the stereotaxic positioning of the recording electrodes on the frontoparietal skull. The deep narco-analgesia lasted about 2.5 h, the time needed to complete all the surgical procedures.

### Pentobarbital-induced sedation

At the end of the surgery, the body temperature was set to and maintained at 37.2°C. The analgesic pentobarbital-induced sedation (light narco-analgesia) was initiated about 2 h after the induction of the surgical narco-analgesia (Figure 1A) and was maintained by a continuous intravenous infusion of the following regimen (average quantity given per kg and per hour): Pentobarbital (7.2 ± 0.1 mg), fentanyl (2.4 ± 0.2 μg), and glucose (48.7 ± 1.2 mg). In order to help maintain the ventilation stable and to block muscle tone and tremors, a neuromuscular blocking agent was used (d-tubocurarine chloride: 0.64 ± 0.04 mg/kg/h). The cortical EEG and heart rate were under continuous monitoring to adjust, when necessary, the infusion rate to maintain the sedation. The EEG recordings began 2 hours after the beginning of the infusion of the sedative regimen. During the recording session and every 2 hours, drops of the local anesthetic lidocaine were applied to the surgical wounds.

**Figure 1.**
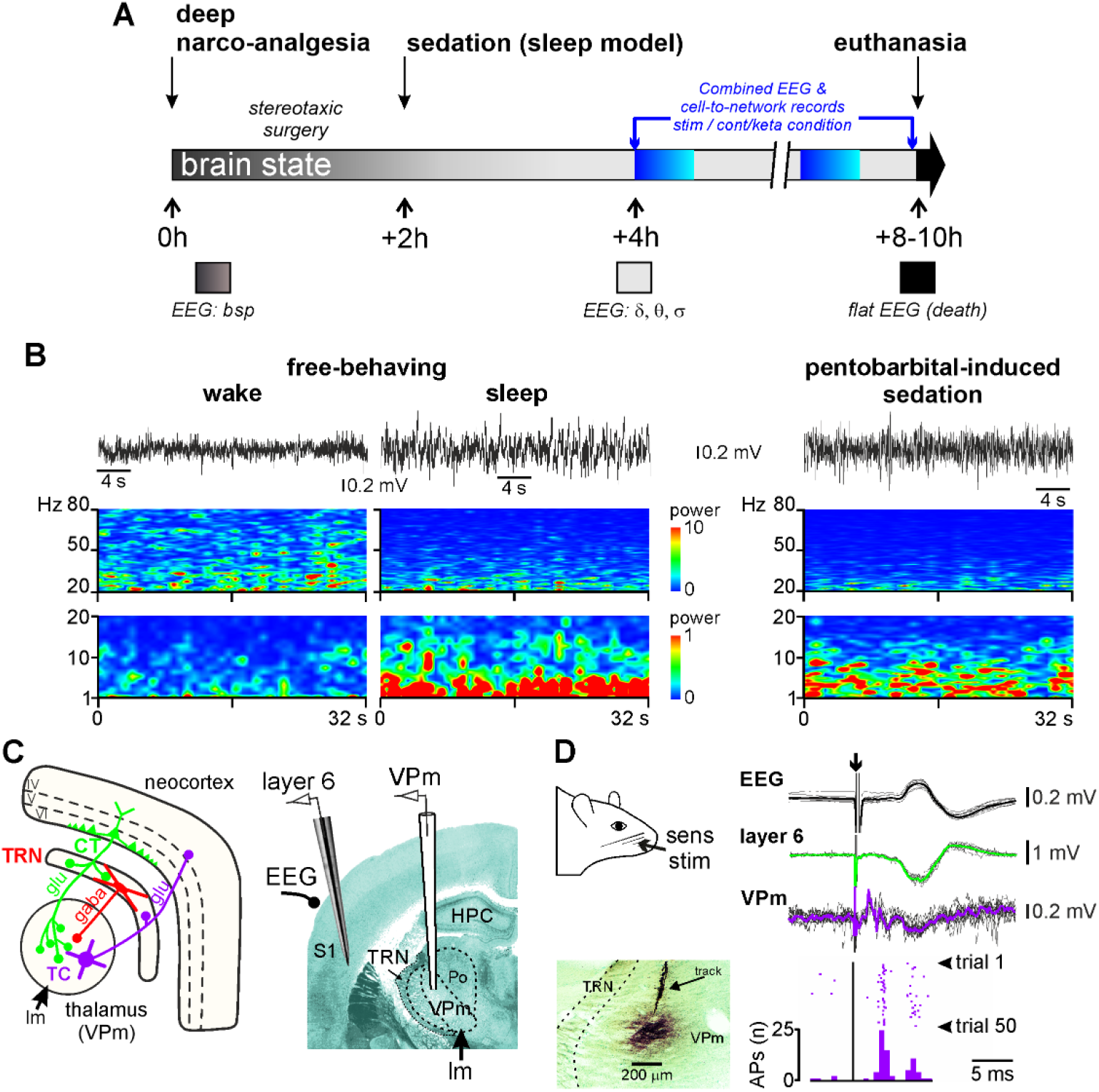
Experimental design. **(A)** Timeline illustrating the key events during the experimental procedure with repeated measures in one animal. One to three low-dose ketamine challenges can be done during one animal-experiment. At the bottom, the color code of the brain state is dark gray for deep narco-analgesia, light gray for sedation (light narco-analgesia), and dark for death. During deep narco-analgesia, the EEG is characterized by a Burst Suppression Pattern (bsp) and, during the sedation, principally by delta-(1-4 Hz), theta-(5-9 Hz) and sigma-(10-17 Hz) frequency oscillations (δ, θ, and σ, respectively). **(B)** Cortical EEG oscillations in the free-behaving (left panel) or pentobarbital sedated (right) rat. Top : 32-s bouts of desynchronized, during wake state, and synchronized, during non-REM sleep, cortical EEG recorded in a free-behaving rat during a 90-minutes recording session; the right trace is from a pentobarbital-sedated rat. Bottom: Time-frequency spectral analysis (resolution: 0.03 Hz, hamming, 50% overlap) of a 32-s recording episode for each condition. The power scale (z scale in color) is not the same for the two frequency bands 1-20 Hz (x1) and 20-80 Hz (x10). The records from the free-behaving rat are from the study performed by Pinault (Biol Psychiatry, 2008). **(C)** Simplified hodology of the thalamocortical (TC, in blue purple) and corticothalamic (CT, in green) pathways of the somatosensory system linked to the vibrissae. This involves the inhibitory afferents originating from the thalamic reticular nucleus (TRN, red). CT and TC neurons are glutamatergic and TRN neurons GABAergic. The right panel shows multi-site recordings within the thalamus (VPm) and neocortex (layer VI and cortical EEG) within the rat somatosensory system. The VPm receives lemniscal afferents (lm). **(D)** The teguments of the vibrissae are electrically stimulated every 15 seconds (sens stim). Sensory evoked potentials are recorded simultaneously within the cortex and the thalamus. For each recording site are shown an overlay of 15 recordings and their averaging. Extracellular field potentials can be accompanied by cellular discharges. The action potentials were identified using the spike sorting methods. In this example, the number of action potentials from the recordings within the VPm are shown (50 trials). At the end of the recording session, the location of the recorded neurons are labelled (extracellular iontophoresis) with the neuronal tracer Neurobiotin. The microphotograph shows the corresponding extracellular labelling (methyl green counterstaining) of the recording site and electrode track in the VPm,. HPC: Hippocampus; Po: Posterior nucleus of the thalamus.

Under the pentobarbital-induced slow-wave sleep (ketamine-free) condition, the EEG recordings principally displayed oscillations in the delta-frequency band (1–4 Hz or slow-waves) accompanied by oscillations in the sigma band (10–17 Hz or “spindle-like” activities) (Pinault *et al*., 2006; Mahdavi *et al*., 2020). These oscillations were qualitatively similar to slow-wave sleep with spindles recorded in free-behaving rats in stage II sleep (Figure 1B). The slow-wave sleep-type oscillations were sometimes interspersed with smaller and faster oscillations, including, among others, broadband gamma- and higher-frequency oscillations.

### Electrophysiology-anatomy

For the EEG recordings of the frontoparietal somatosensory cortex (stereotaxic coordinates relative to bregma (Paxinos & Watson, 1998): posterior 2.3 mm, lateral 5 mm), the section of the Ag/AgCl wires (diameter 150 μm), insulated with Teflon, was placed on the inner plate of the bone. Extracellular field potential and multi-unit activities were recorded in the somatosensory thalamus, especially in the medial part of the ventral posterior nucleus (VPm, bregma −2.8 mm, lateral 2.8 mm), and in the medial part of the posterior group (PoM, bregma −2.8 mm, lateral 2.8 mm, depth 5.6 mm.) using glass micropipettes (tip diameter of 5-10 μm) filled with artificial cerebrospinal fluid and 1.5 % Neurobiotin). Semi-micro quartz/platinum-iridium electrodes were used for recordings in the layer 6 (depth 1.6-2.0 mm) of the related somatosensory cortex (Figure 1C). All regions of interest were recorded simultaneously, and the electrophysiological signals were sampled with a rate of 20 kHz (Digidata 1440A with pCLAMP 10 Software, Molecular Devices). The recording electrodes were moved down until the electrophysiological identification of the receptive field (Figure 1D). The anatomical identification of the recording site was validated following extracellular iontophoresis of the neuronal tracer Neurobiotin (Figure 1D), which was revealed using a standard histological procedure (Pinault, 1996). Sensory-evoked potentials were recorded after electrical stimulation of the vibrissae teguments using a pair of subcutaneous needles (duration: 75 μs; intensity: 50–60% of the intensity that gives maximal amplitude evoked potential, 1.0 to 1.5 mA; frequency: 0.06 Hz). Every trial contained recorded signals after one stimulation (10 trials/rat).

### Pharmacology and repeated measures in one animal

During the recording session under the sedation condition, every rat was its own control. Saline (vehicle, NaCl 0.9%) and ketamine (2.5 mg/kg) were subcutaneously administered (1ml/kg). As long as the pentobarbital-induced sedation is stable (4 to 6 hours) and knowing that, under the present experimental conditions, the ketamine effect (peaking at 15–20 min) lasts significantly less than 90 min (Anderson *et al*., 2017; Mahdavi *et al*., 2020), two to three conditions (20-40-minutes saline condition followed by one or two ketamine challenges) could be performed in one animal (Figure 1A).

### Data Analysis

Data analyses were performed with Clampfit 10, SciWorks (Datawave Technologies) and MATLAB softwares. Spectral analysis was done with 2 Hz resolution, hamming windowing, none overlay. Recorded signals were analyzed in five frequency bands: delta (1-4 Hz), theta (5-9 Hz), sigma (10-16 Hz), beta (17-29 Hz), and gamma (30-80 Hz). Sensory stimulus-induced gamma power was calculated by subtracting the evoked and basal power from the total power.

#### Total power spectral

In total, data of 40 trials (10 × 4 rats) were acquired. The total power in a given frequency range *v* was the sum of powers across the defined spectral range. We consider the 500 ms period before each stimulus as baseline. The baseline power for each trial is denoted as *P*_*b1*_, *P*_*b2*_, *P*_*b3*_, *…P*_*b40*_; stimulus-related power for each 200-700ms post-stimulus 500-ms epoch is denoted as *P*_*e1*_, *P*_*e2*_, *P*_*e3*_, *… P*_*e40*_. Therefore, the average normalized power *P*_*v*_ for the frequency *v* range is computed as:

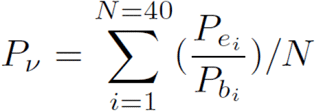

Similarly, in ketamine conditions, *P*_*kbi*_ and *P*_*kei*_ stand for the baseline and stimulus-related power respectively for each trial *i*. Thus, average normalized ketamine power P_kv_ is computed as:

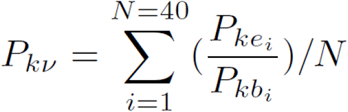

#### Multi-scale entropy (MSE)

The information complexity of extracellular LFP beta- and gamma-frequency oscillations (raw LFP filtered at 17-29 Hz and 30-80 Hz, respectively) was measured by MSE. Although the definition of complexity is various, it is associated with “meaningful structural richness” and “information randomness” (Costa *et al*., 2005; Hager *et al*., 2017). The MSE is calculated with 2 steps. First, coarse-graining is applied to the time series {x_1_, …, x_i_, …, x_N_}. It is constructed by averaging data points from non-overlapping time-windows of interest, τ. Every coarse-grained time series, *y*_j_^τ^, is calculated as :

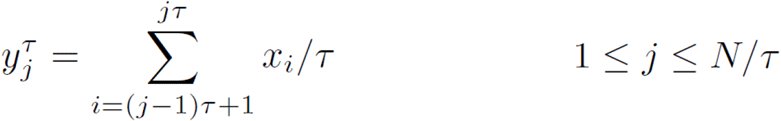

Where *N/τ* is the length of each resulting coarse-grained time series. Then the sample entropy is calculated for each series *y*_j_^τ^ and plotted as function of the scale factor. When τ equals one, *y*_j_^1^ is equivalent to the original time series. The higher scale factor is, the longer temporal range it is. The MSE values for low scales reflect short-range temporal irregularity, while high scales reflect long-range temporal irregularity. Other parameters for MSE calculations were adopted from previous studies (Lake *et al*., 2002; Richman *et al*., 2004; Takahashi *et al*., 2010).

#### Coherence connectivity

Coherence was calculated by the MATLAB mscohere function.

#### Statistics

All statistics tests were calculated using MATLAB or Graphpad Prism 9. For comparing the baseline and stimulus-related activities in different frequency bands, we used one-way ANOVA test with Holm-Šidák’s multiple comparisons test as post-hoc analysis. For comparing normalized gamma total power under ketamine and saline conditions, paired t-tests were applied to each group, each animal being its own control. For testing the effect of ketamine on induced gamma oscillations, we computed whether there was a significant interaction using a two-way ANOVA for time and condition (saline or ketamine). When assessing the coherence between recording sites, we used the Wilcoxon matched pairs signed-rank test.

## RESULTS

### Ketamine increases baseline and decreases induced beta/gamma oscillations

A representative example of multisite cortical (EEG and layer 6) and thalamic (VPm and PoM) recordings, filtered at the beta/gamma frequency band, is shown in figure 2. It reveals the ongoing (baseline) and sensory-related oscillations 2-s before and 2-s after the stimulation (0 s), respectively. It is striking that ketamine increased the power of ongoing beta/gamma oscillations in the cortex and thalamus (Fig. 3 A and B) as demonstrated in previous studies (Pinault, 2008; Hakami et al, 2009; Kocsis, 2012). Immediately and later after the sensory stimulation, the amplitude of the beta/gamma oscillations was modulated. During the 200-700 ms post-stimulus period, the induced beta/gamma oscillations significantly decreased in power at all recorded cortical and thalamic sites following ketamine administration (Fig. 3C and D).

**Figure 2.**
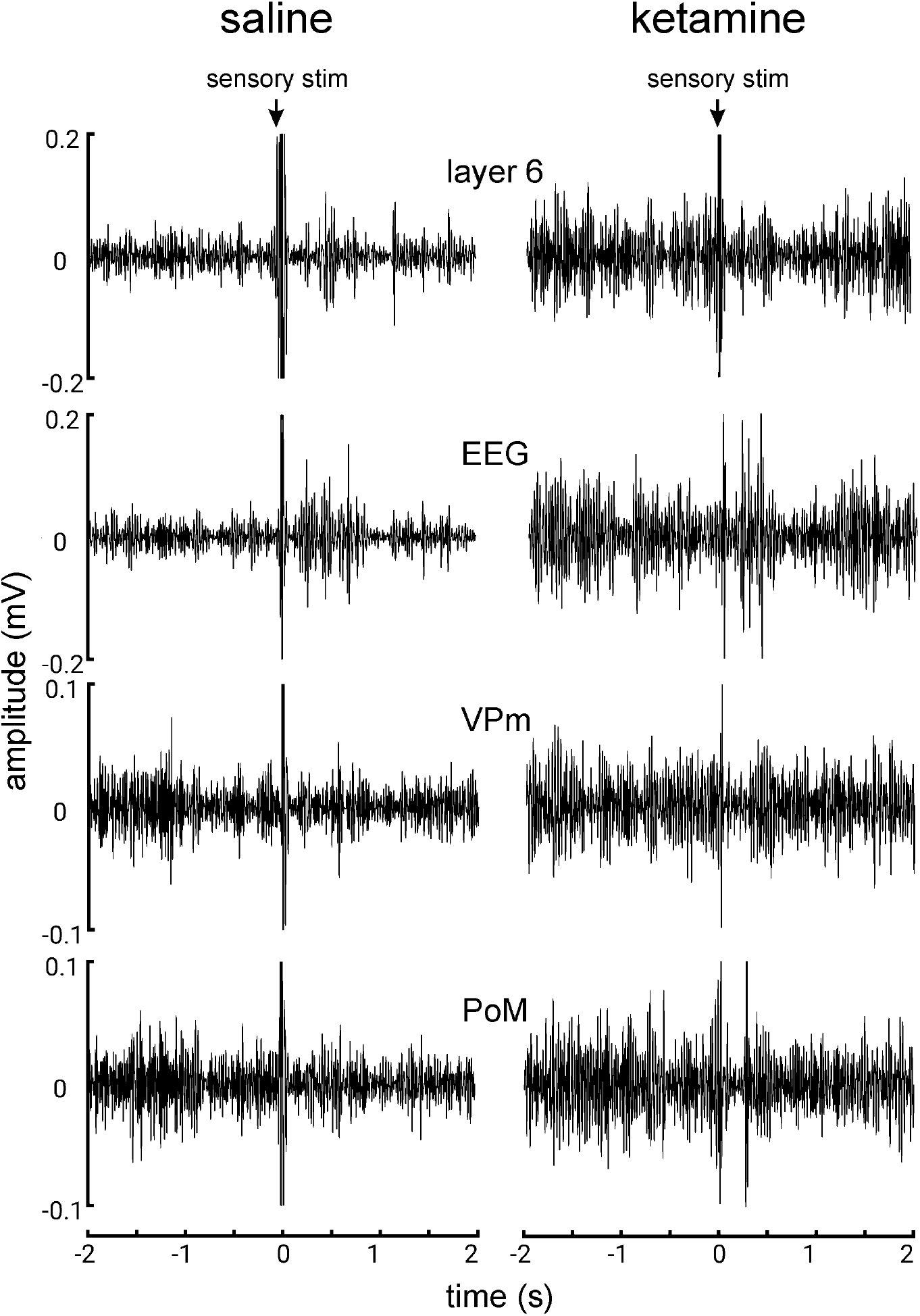
Simultaneous cortical (EEG & layer 6) and thalamic (VPm & PoM) recordings under saline and ketamine conditions from a lightly anesthetized rat. They are filtered in the beta/gamma frequency band (25-50 Hz). Each trace (sweep) shows the 2-s pre-stimulus and 2-s post-stimulus periods. The teguments of the vibrissae are stimulated (sensory stim). The traces under the saline condition were recorded 10 minutes before the ketamine administration. The ketamine traces were recorded 20 minutes after the subcutaneous administration of ketamine at a subanesthetic low-dose (2.5 mg/kg).

**Figure 3.**
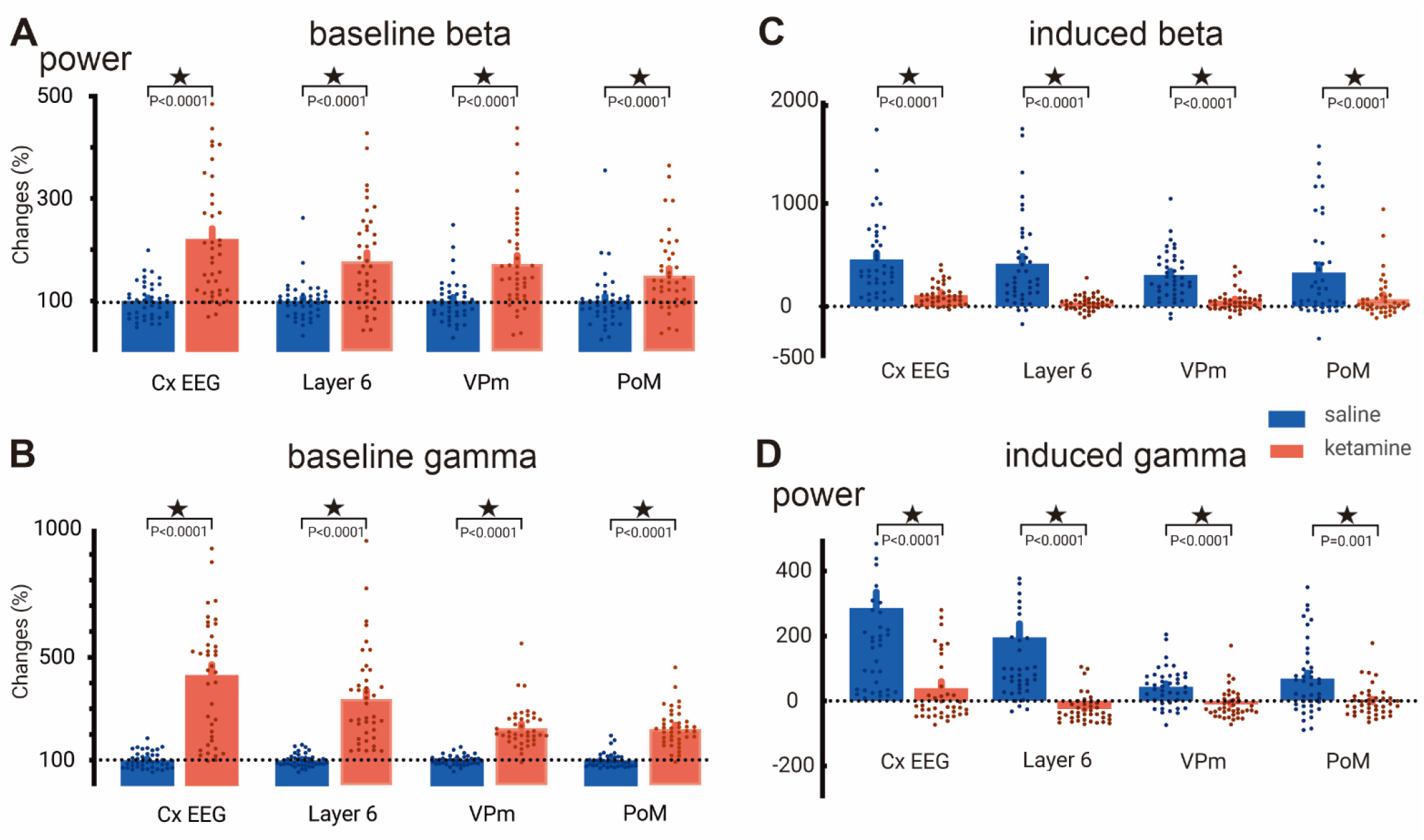
Ketamine increases baseline beta/gamma (A/B) oscillations, and decreases sensory-induced beta/gamma (C/D) oscillations. **(A, B)** Each column stands for the average (± SEM, from 40 values, 10 per rats, 4 rats) of the normalized total power of gamma oscillations relative to saline baseline. Asterisks when significant (paired t-test, all P value < 0.0001, check supplement for statistic details). **(C, D)** The power of the induced gamma oscillations was obtained when subtracting the power of the baseline gamma from the power of the sensory-elicited total gamma. Each value (± SEM, from 40 values, 10 per rats, 4 rats) is the % change relative to the baseline gamma recorded before the sensory stimulation. Asterisks when significant (< 0.05, two-way ANOVA, corrected with Holm-Sidak, check supplement for details).

### Ketamine disrupts information transferability in the corticothalamic (CT) network

The post-stimulation time lapse of 200-700 ms is long enough to encode, integrate, and perceive the incoming sensory signal (Rodriguez *et al*., 1999). We hypothesized that ketamine can interrupt information processing during this period. To test this, we measured the uncertainty of information based on MSE in the gamma band during the 200-700 ms post-stimulus period (Figure 4). The measurement of entropy can be used as an estimation of “complexity” in physiological systems. Higher entropy means the system is likely in a more “complex/dynamic” state (Costa *et al*., 2005; Hager *et al*., 2017). In figure 4, it is shown that the entropy of the VPm and that of layer 6 were significantly increased along different time scales.

**Figure 4.**
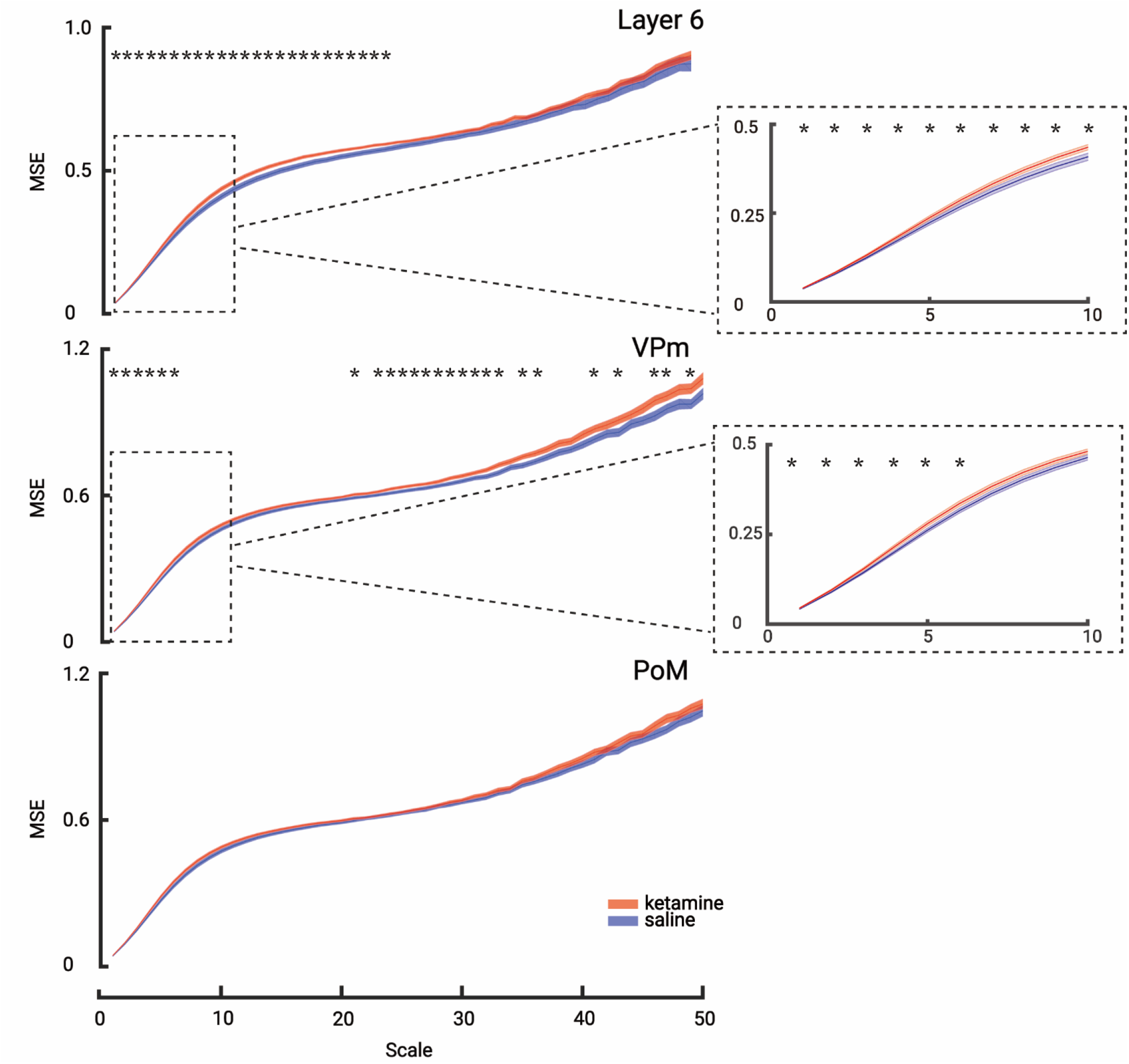
Ketamine increases multi-scale entropy in the gamma band between the VPm and layer 6. Comparison of the multi-scales entropies of saline (blue) and ketamine (red) conditions at all recording sites in the gamma band during the 200-700 ms post-stimulus period. Layer 6 and VPm show a significant increase in entropy. Each scale point is the average entropy (± SEM, from 40 values, 10 per rats, 4 rats). Asterisks when significant (< 0.05, paired t-test). Mean values: Layer 6: t(39) = 2.5509 (p<0.05); VPm: t(39) = 2.2930 (p<0.05).

This indicates that the information contained in the VPm and layer 6 extracellular potential was biased toward a random or “noisy” state. No difference was observed in the PoM (Paired t-test, p<0.05). Also, no significant difference was observed on MSE for the beta band in both the cortex (layer 6) and the thalamus (VPm and PoM) (Figure S1).

The present study shows that ketamine had a strong effect on information in the specific thalamic nucleus (VPm) and in the related layer 6 in the gamma band, indicating that the gamma connectivity in the whole network might also be dysfunctional. So, we used the coherence coefficient to measure the induced connectivity in the beta and gamma bands between the cortex and thalamus. We applied the magnitude-squared coherence to measure similarities between two signals during the 200-700 ms post-stimulus period. Ketamine elicited a non-significant decrease (∼ 25%) of the coherence coefficient between layer 6 and the related thalamus (VPm) in the beta band connectivity (Fig. 5A). On the other hand, in the gamma band connectivity, ketamine dramatically decreased (nearly 40 %) the coherence between layer 6 and the VPm (Fig. 5B). Furthermore, no significant change of coherence was observed between PoM and VPm, or layer 6 and PoM in both the beta and gamma connectivity (Figure 5A,B, Wilcoxon matched-pairs signed-rank test).

**Figure 5.**
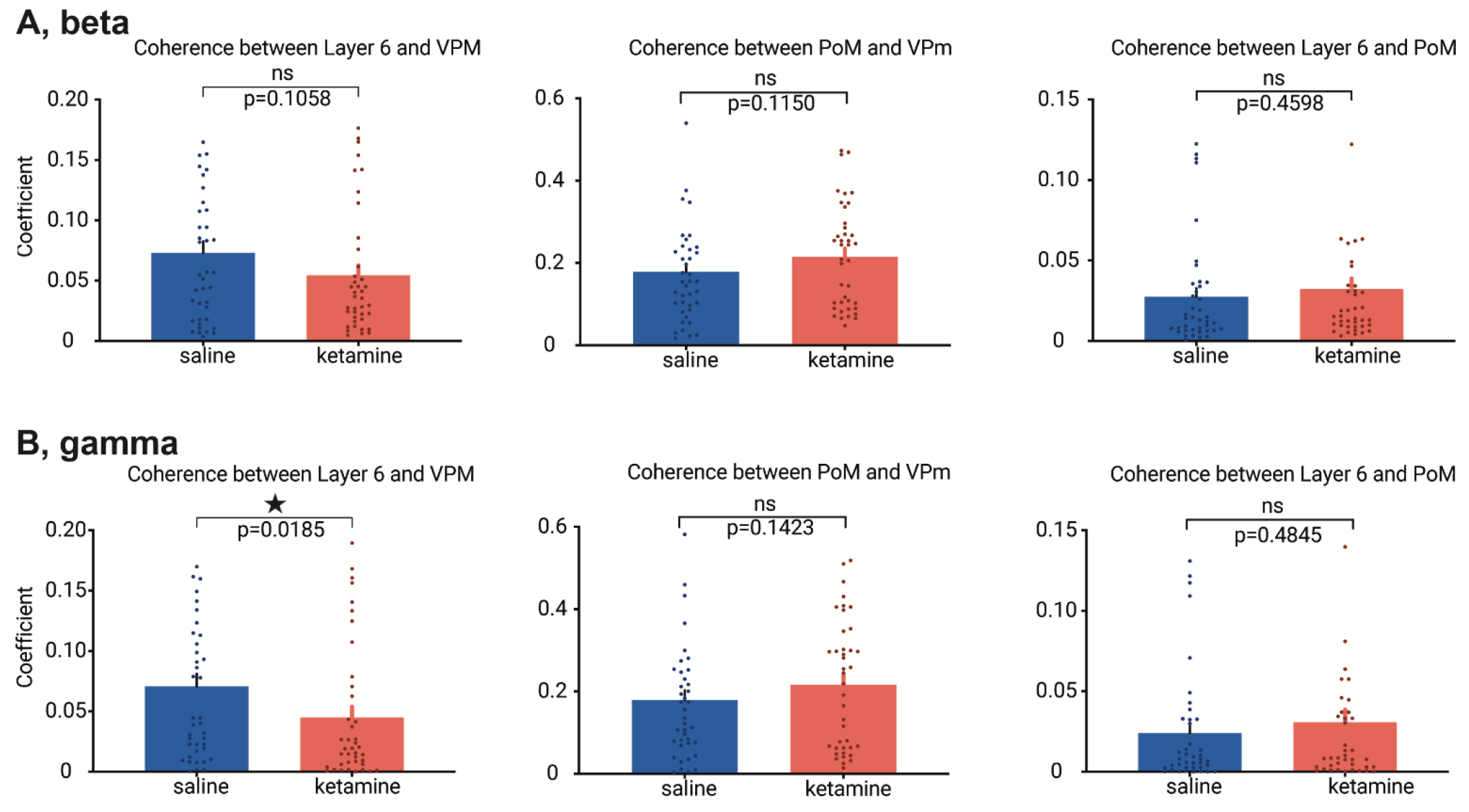
Ketamine decreases coherence connectivity in the beta/gamma band between layer 6 and VPm. Comparison of coherences connectivity of beta/gamma bands within CT-TC network under saline and ketamine conditions. Blue and red columns stand for the average coefficients (± SEM, from 40 values, 10 per rats, 4 rats) of the coherences of beta (A)/gamma (B) oscillation in the saline and ketamine conditions, respectively. Asterisks when significant (< 0.05), Wilcoxon test.

## DISCUSSION

The findings presented here demonstrate that the parietal CTC system significantly contributes to sensory stimulus-induced thalamic beta/gamma frequency oscillations, which occur at the late post-stimulus stage (200-700 ms), and that the administration of the NMDA receptor antagonist ketamine disrupts the transfer of perceptual information in the system.

### Low-dose ketamine decreases the signal-to-noise ratio

The present study analyzed beta (17-29 Hz) and gamma (30-80 Hz) oscillations separately. Following the subcutaneous administration of ketamine, spontaneously occurring beta and gamma oscillations increased in amplitude and power in the CT system. Furthermore, the sensory-induced beta and gamma oscillations were significantly decreased in power at all recorded cortical and thalamic sites; ketamine increased the MSE in the specific CT system (Layer 6-VPm) in the gamma but not in the beta frequency band; ketamine decreased the coherence between layer 6 and VPm in both the beta (not significant) and gamma (significant) frequencies. These findings suggest that beta and gamma oscillations had a common functionality, which is supported by a previous study demonstrating that, in the cerebral cortex, both frequency bands share cellular and biophysical mechanisms (Compte *et al*., 2008). Along these lines, intracellular recordings of TRN neurons demonstrated that the frequency of the intrinsically generated membrane potential oscillations responsible for generating gamma frequency (25-60 Hz) firing can drop up to 18 Hz (Pinault & Deschenes, 1992).

In the lightly-anesthetized rat, vibrissae stimulation generates a wide-band frequency response in extracellular recordings simultaneously in the VPm, PoM, and in the related somatosensory cortex (layer 6). Sensory-induced beta- and gamma-frequency oscillations were significantly decreased in a smaller post-stimulus time window (200-700 ms). The ketamine-induced obliteration of the sensory stimulus-induced beta/gamma oscillations at 200-700 ms may be the result of the ketamine-elicited abnormally and diffusely amplified basal gamma oscillations, which lead to disruption of beta/gamma-related information transferability in the somatosensory CTC system as assessed, in the present study, by an increase in MSE and a decrease in coherence connectivity in the specific CT system.

The fact that ketamine aberrantly and diffusely amplifies ongoing beta/gamma oscillations at all the recorded cortical and thalamic sites supports a conception of NMDA receptor hypofunction-related beta and gamma hyper-synchronies as an aberrant generalized diffuse network noise. This NMDA receptor hypofunction would induce a noise state then would contribute to disrupting the ability of neural networks to encode and integrate input signals. In other words, NMDA receptor antagonism decreases the signal-to-noise ratio (Hakami *et al*., 2009; Gandal *et al*., 2012; Kulikova *et al*., 2012; Anderson *et al*., 2017). The disruption of the transfer of sensory information would start to occur at least during the very first stages (up to ∼ 15 ms) of information processing, at the gate of cognitive processes (Briggs & Usrey, 2008; Kulikova *et al*., 2012; Anderson *et al*., 2017; Homma *et al*., 2017). The aberrant diffuse network beta/gamma noise may be a potential electrophysiological correlate of a psychosis-relevant state as increased gamma synchrony has been recorded in patients during somatic and visual hallucinations (Baldeweg *et al*., 1998; Behrendt, 2003; Spencer *et al*., 2004; Becker *et al*., 2009) and, importantly, in clinically at-risk mental state patients for psychosis transition and naïve in antipsychotic medication (Ramyead *et al*., 2015; Perrottelli *et al*., 2021).

### Ketamine-induced increase in randomness or complexity in a signal

Our MSE results show that the specific thalamic nucleus (VPm) and related layer 6 somatosensory cortex have increased sample entropy (i.e. complexity) following ketamine administration, which is consistent with previous studies on human patients with schizophrenia (Takahashi *et al*., 2010). Of particular interest is that both layer 6 and the VPm showed increased MSE in the lower time scale factors, meaning those with the most detailed temporal information (i.e., with more high-frequency information incorporated in the complexity measure at these time scales). Interestingly, there is a tendency for younger, medication-free patients with higher positive symptoms to display higher levels of complexity in EEG (Fernandez *et al*., 2013), a finding that matches our interpretation of ketamine administration to model a psychotic-like state. It has further been suggested that these increases in neural complexity measures are evidence for the “disconnection hypothesis” (Friston *et al*., 2016) whereby disruption (aberrant or reduced) in connectivity increases EEG signal complexity (Takahashi *et al*., 2010). In healthy humans submitted to a cognitive-visual task, ketamine increases the power of broadband gamma oscillations and disrupted feedforward and feedback signaling, leading to hypo- and hyper-connectivity in CTC networks (Grent-’t-Jong *et al*., 2018).

Increased entropy may also be interpreted as an increase in “randomness” in a signal (with a truly random signal having maximal entropy (Ahmed & Mandic, 2011). Applying such an interpretation to our present results would reflect ketamine administration adding “noise” to the system increasing ongoing beta/gamma power, resulting in a more random signal. The increased ongoing gamma power reflects an aberrant and pathological increase in non-relevant beta/gamma activity that effectively attenuates sensory-related induced beta/gamma interfering with its sensory transmission. This decreases the overall beta/gamma signal-to-noise ratio in the CTC system, disconnecting these areas and impairing sensory perception. Accompanying this disconnection hypothesis, we also observed a functional disconnection of phase coherence measures in the gamma frequency band between layer 6 and VPm, but not between PoM and layer 6 or VPm. This result supports the interpretation that sensory-generated gamma activity has been disrupted between cortex and thalamus under the ketamine condition. Disorders in the intrinsic properties (amplitude, noise, and complexity) and spatial dynamics (coherence) of gamma oscillations somehow reflect a fundamental disturbance of basic integrated brain network activities.

### Possible underlying mechanisms

The CTC system is involved in multiple integrative functions, including sensory, perceptual, and attentional processing (Pinault, 2004; Van Essen, 2005; Wolff *et al*., 2021). Sensory-to-perceptual responses of the CTC system result from dynamic interactions between TC (bottom-up) and CT (top-down) processing (Alitto & Usrey, 2003; Briggs & Usrey, 2008; Homma *et al*., 2017). Both TC and CT pathways are glutamatergic. In thalamic neurons, the response pattern depends on the brain state (Castro-Alamancos, 2002; Urbain *et al*., 2015) and on thalamic GABAergic inhibition that is mediated principally by the external source thalamic reticular nucleus (TRN) (Pinault, 2004). Under the present experimental sleep conditions, almost all the thalamic glutamatergic and GABAergic neurons are hyperpolarized and fire in the burst mode (Pinault *et al*., 2006; Mahdavi *et al*., 2020), and the arousal promoting effect of ketamine switches the spontaneous firing pattern of both the glutamatergic TC and the GABAergic TRN neurons from the burst to the tonic mode (Mahdavi *et al*., 2020).

The post-inhibitory rebound excitation is a cellular intrinsic property, which occurs during physiological and pathological brain oscillations or following the activation of prethalamic (e.g., sensory) or cortical inputs. For instance, during slow-wave sleep, a long-lasting hyperpolarization gives rise, in the thalamic relay and reticular neurons, to a rebound excitation caused by a low-threshold calcium-dependent potential, de-inactivated by membrane hyperpolarization, and can be topped by a high-frequency burst of action potentials (Deschenes *et al*., 1984; Jahnsen & Llinas, 1984; Llinas, 1988; Grenier *et al*., 1998; Urbain *et al*., 2019). Such a post-inhibitory rebound excitation is also recorded under anesthesia in TC neurons following the activation of prethalamic or cortical inputs (Deschenes *et al*., 1984; Grenier *et al*., 1998). There is evidence that TC bursting may serve as a “wake-up call” in the initiation of perceptual/attentional processes (Sherman, 2001; Swadlow & Gusev, 2001). Two potential mechanisms of the effect of the NMDA receptor antagonist ketamine could be responsible alone or in combination: 1) reduced TRN-mediated inhibition (see discussion by Mahdavi et al, 2020), and 2) a reduction of the hyperpolarization-activated cationic current Ih (Kim & Johnston, 2020). Of course, ketamine also acts at multiple cortical and subcortical structures, and there is increasing evidence that it suppresses the activity of GABAergic interneurons leading to disinhibition of glutamatergic neurons (Homayoun & Moghaddam, 2007; Ali *et al*., 2020). Although ketamine acts at many other receptors, including dopaminergic, serotoninergic, opioid, and GABAergic receptors (Sarton *et al*., 2001; Kapur & Seeman, 2002; Seeman & Lasaga, 2005; Lewis *et al*., 2008), it is generally believed that most of its effects are accounted for by NMDA receptor antagonism.

The late (200-700 ms post-stimulus) sensory-induced beta/gamma oscillations were, on average, measurable in the sedated rat. These late response activities would be involved in the perceptual process (Funayama *et al*., 2015). In sedated rats, the level of perception and the underlying neural activities are expected to be attenuated because of the presence of rhythmic GABAergic mediated inhibitions at least in the delta- and sigma-frequency bands (slow-wave sleep with spindles). Ketamine, by reducing the slow waves, spindles, and burst activities, depolarizes and switches the burst firing pattern to the irregular tonic mode (Mahdavi *et al*., 2020). This means that ketamine brings a persistent depolarizing pressure to the membrane potential (persistent UP state) of the glutamatergic and GABAergic neurons, which is expected to disrupt the tonic firing pattern associated with a sensory-perceptual process. Moreover, it was demonstrated that, in the rat, NMDA receptor antagonism disrupts synchronization of action potential firing in the prefrontal cortex, which would lead to a disruption of the transfer of information processing dependent on the timing of action potentials (Molina *et al*., 2014).

### Limitations of the study

As the experiments were performed in the pentobarbital-sedated rat (non-REM sleep model), the present findings do not allow drawing definitive conclusions. However, the combined two models, one for the brain state (sleep model) and one for the ketamine psychosis-relevant challenge (Mahdavi *et al*., 2020), provide interesting tools to conceptually and mechanistically advance in the understanding of the neurobiology of psychotic disorders. The major requirement of the present study was to have the equivalent of a stationary stage II non-REM sleep, during which we could perform repeated measures. Furthermore, under the present experimental conditions, a single systemic administration of ketamine at a psychotomimetic dose (2.5 mg/kg, estimated from a study conducted in free-behaving rats (Pinault, 2008) induces most of the oscillopathies (especially basal gamma hyperactivity and delta/spindle hypoactivity) recorded in patients having psychotic disorders (Mahdavi *et al*., 2020). One advantage of the pentobarbital-induced sedation was its relative stability over time allowing repeated measures.

Under the present experimental conditions, the degree of perception and attention would have been weakened because the pentobarbital induces a slow-wave sleep with spindle-like activities by increasing the GABAergic neurotransmission (Maldifassi *et al*., 2016). In short, the experimental conditions, which promote cortical slow waves (Pinault *et al*., 2006; Murphy *et al*., 2009; Knyazev, 2012; Urbain *et al*., 2019), would have prevented or attenuated the full corticalization of sensory-perception processing and, subsequently the full implication of the CT (feedback) and cortico-cortical (feedforward) pathways. However, the CTC system, which includes the TRN (Pinault, 2004), is functionally polyvalent in a state-dependent way as it is involved in sensory-perception processing (Saalmann & Kastner, 2011), the wake-sleep brain oscillations (Steriade *et al*., 1993), and in attention-sensory processes (Chen *et al*., 2015; Wimmer *et al*., 2015). So, it is not surprising that the highly distributed CTC systems play a central role in the disorders of sleep integrity, sensorimotor, perception, and attentional processes observed in patients with psychotic disorders (Steriade *et al*., 1993; Shipp, 2004; Ferrarelli & Tononi, 2011; Pinault, 2011; Chen *et al*., 2015; Wolff *et al*., 2021).

Ketamine exerts a wide range of dose-dependent effects (dissociative anesthesia, sedation, psychotomimetic, antidepressant, analgesic…). So, we should not exclude the analgesic effect of low-dose ketamine (Laskowski *et al*., 2011; Zanos *et al*., 2018). Pain involves the somatosensory CTC system among many other brain regions (amygdala, insula, S2, ACC, and PFC). By examining the role of the auditory system in pain processing, it was demonstrated that the information to VPm and PoM is disrupted by the auditory CT pathway (Zhou *et al*., 2022). So, the ketamine-induced disruption of sensory information transfer in the CT network may be a common part of the mechanisms underlying the analgesic and psychotomimetic effects of ketamine.

### Conclusion and perspectives

The present results provide anatomo-functional relevance to understanding the neural dynamics underlying ketamine-induced impairment of encoding processes (Hetem *et al*., 2000), perception-related (feedforward and feedback) dysconnectivity, and abnormal amplification of gamma oscillations in human CTC systems (Driesen *et al*., 2013; Anticevic *et al*., 2014; Hoflich *et al*., 2015; Rivolta *et al*., 2015; Grent-‘t-Jong *et al*., 2018). The NMDA receptor hypofunction-related gamma hyper-synchronies (power increases) are neurophysiological abnormalities that may represent a core biological feature of the psychotic transition. Although the interpretation of measures using complexity estimators (like MSE) of neural signals is not simple, in recent years there is accumulating evidence that increased and abnormal complexity may also be a hallmark of psychosis (Fernandez *et al*., 2013; Yang *et al*., 2015; Ibanez-Molina *et al*., 2018). Abnormal and diffuse amplification of spontaneously-occurring broadband gamma oscillations in neural networks (gamma noise) associated with reductions in sensory-related, evoked, and induced gamma-band responses (gamma signal) are potentially predictive translational biomarkers of psychosis transition (Hakami *et al*., 2009; Gandal *et al*., 2012; Kulikova *et al*., 2012; Anderson *et al*., 2017). The sensory-evoked potential is also an appropriate index to evaluate the expression of the plasticity of neural circuits (Kulikova *et al*., 2012). Because of their spatio-temporal structure and stereotyped pattern, sensory-evoked and induced gamma oscillations represent potential reliable and suitable variables (Spencer *et al*., 2008; Hong *et al*., 2010; Leicht *et al*., 2016; Tada *et al*., 2016; Reilly *et al*., 2018) for the development of innovative therapies preventing the psychotic transition.

## LIST OF ABBREVIATIONS

CT: corticothalamic
CTC: cortico-thalamo-cortical
EEG: electroencephalography
GABA: gamma-aminobutyric acid
LFP: local field potential
MSE: multiscale entropy
NMDA: N-methyl-D-aspartate
PoM: medial part of the posterior group
TC: thalamocortical
TRN: thalamic reticular nucleus
VPm: medial part of the ventral posterior nucleus

## ACKNOWLEDGEMENTS

The present work was supported by INSERM, the French National Institute of Health and Medical Research (Institut National de la Santé et de la Recherche Médicale, 2013-), l’Université de Strasbourg, Unistra (2013-), and Neurex. This project has been funded with support from the NeuroTime Erasmus+ program of the European Commission (2015-2020: YQ and AM). This publication reflects the views only of the authors, and the Commission cannot be held responsible for any use, which may be made of the information contained therein. The authors thank Damaris Cornec for her technical assistance, and Christiaan Levelt for critical reading of the manuscript.

## DISCLOSURES

All authors have approved the final version of the article. The authors report no competing biomedical financial interests or potential conflicts of interest.

## Notes

### Competing Interest Statement

The authors have declared no competing interest.

### Summary of Updates

.Abstract: reworded .Introduction: The first para rewritten .Methods/Pentobarbital-induced sedation: Experimental conditions, 2nd para (moved from 1st draft results). .Methods/Electrophysiology-anatomy: Justification on the use of the neuronal tracer Neurobiotin .Methods/Data analysis/MSE: Precision on LFP .The presentation of the data has been amended and simplified. .Discussion: the 2nd para on beta and gamma oscillations has been added. .Discussion/Limitations of the study, last para on the potential mechanisms underlying the analgesic effects of low-dose ketamine. The mechanisms may be similar to those of the psychotomimetic effect of ketamine. .Supplemented files updated.

