## Supplemental Figure 1 for "The psychotomimetic ketamine disrupts the transfer of late sensory information in the corticothalamic network"

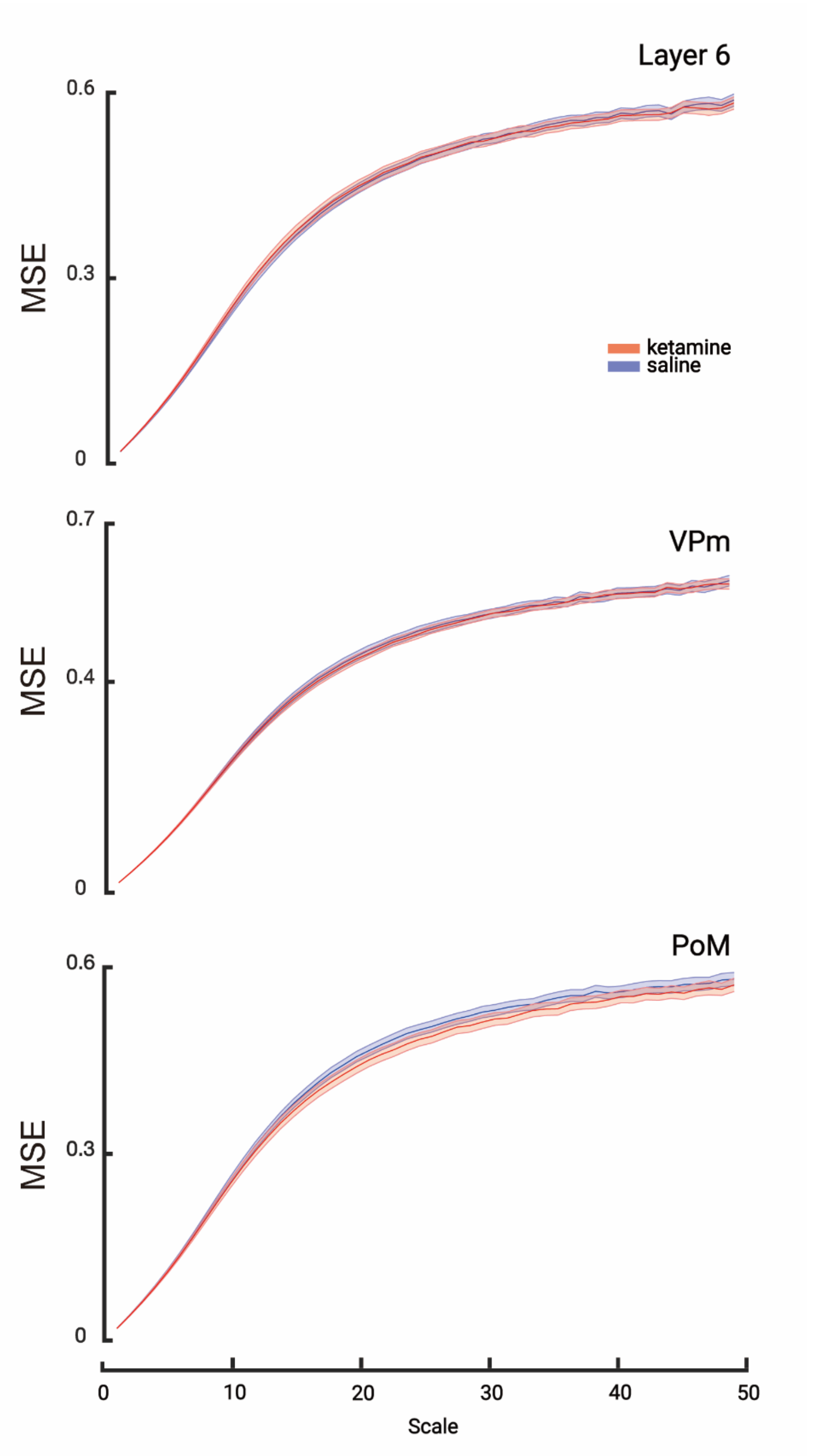


**Figure S1: Ketamine does not change the multi-scale entropy in the beta band between the cortex and thalamus.** Comparison of the multi-scales entropies of saline (blue) and ketamine (red) conditions at all recording sites in the beta band during the 200-700 ms post-stimulus period. Layer 6 and VPm show a significant increase in entropy. Each scale point is the average entropy (± SEM, from 40 values, 10 per rats, 4 rats). The statistical test does not reveal any significant difference between the saline and ketamine conditions ($p > 0.05$, paired t-test).
